# An ATM/Wip1-dependent timer controls the minimal duration of a DNA-damage mediated cell cycle arrest

**DOI:** 10.1101/042119

**Authors:** Himjyot Jaiswal, Jan Benada, Erik Müllers, Karen Akopyan, Kamila Burdova, Tobias Koolmeister, Thomas Helleday, René H Medema, Libor Macurek, Arne Lindqvist

**Affiliations:** Department of Cell and Molecular Biology, Karolinska Institutet, Sweden; Laboratory of Cancer Cell Biology, Institute of Molecular Genetics, Academy of Sciences of the Czech Republic, Czech Republic; Faculty of Science, Charles University in Prague, Czech Republic; Department of Medical Biochemistry and Biophysics, and Science for Life Laboratory, Karolinska Institutet, Sweden; Division of Cell Biology, Netherlands Cancer Institute, the Netherlands; Present address: Discovery Science, Astrazeneca R&D, Mölndal, Sweden

## Abstract

After DNA damage, the cell cycle is arrested to avoid propagation of mutations. In G2 phase, the arrest is initiated by ATM/ATR-dependent signalling that blocks mitosis-promoting kinases as Plk1. Interestingly, Plk1 can counteract ATR-dependent signalling and is required for eventual resumption of the cell cycle. However, what determines when Plk1 activity can resume remains unclear. Here we use FRET-based reporters to show that a global spread of ATM activity on chromatin and phosphorylation of targets including Kap1 control Plk1 re-activation. These phosphorylations are rapidly counteracted by the chromatin-bound phosphatase Wip1, allowing a cell cycle restart despite persistent ATM activity present at DNA lesions. Combining experimental data and mathematical modelling we propose that the minimal duration of a cell cycle arrest is controlled by a timer. Our model shows how cell cycle re-start can occur before completion of DNA repair and suggests a mechanism for checkpoint adaptation in human cells.

## Introduction

DNA double-strand breaks (DSBs) represent a serious threat to the genome integrity of a cell. Failure to recognize and repair these lesions can lead to mutations, genome instability and cancer (Jackson and Bartek, 2009). To cope with DSBs cells launch a DNA damage response (DDR), involving a network of DNA damage sensors, signal transducers and various effector pathways. Besides orchestrating DNA repair, a central component of DDR is activation of a checkpoint that blocks cell cycle progression (Bartek and Lukas, 2007; Medema and Macurek, 2011). This is particularly important in the G2 phase of the cell cycle, as cell division in the presence of DSBs may lead to aneuploidy and propagation of mutations to progeny. Surprisingly however, a growing body of evidence suggests that a cell cycle block is commonly reversed before all DNA lesions are repaired in both transformed and untransformed cells (Deckbar et al., 2007; Loewer et al., 2013; Syljuasen et al., 2006; Tkacz-Stachowska et al., 2011).

A key unresolved issue therefore is how the duration of a cell-cycle arrest is controlled. Upon recruitment to DSBs, the Ataxia telangiectasia mutated (ATM) kinase initiates a signaling cascade by phosphorylating S/TQ motifs in more than 700 proteins, many of which are central proteins in various branches of the DDR (Matsuoka et al., 2007; Mu et al., 2007; Shiloh and Ziv, 2013). However, although crucial for initiating many of the responses, the role for ATM in maintaining a cell cycle arrest remains unclear, as acute inhibition of ATM after a checkpoint is initiated has limited effect on the efficiency of cell cycle resumption (Kousholt et al., 2012). Rather, ATM and Rad3 related kinase (ATR), activated by repair intermediates of DSBs, is a main controller of checkpoint duration (Sanchez et al., 1997; Shiotani and Zou, 2009).

To enforce a cell cycle arrest, ATM and ATR-dependent signaling inhibits the activities of mitosis promoting kinases as Cyclin dependent kinase 1 (Cdk1), Polo-like kinase 1 (Plk1), and Aurora A (Krystyniak et al., 2006; Lock and Ross, 1990; Smits et al., 2000). In particular, ATM, ATR, and p38 activate Chk2, Chk1, and MK2, respectively, structurally distinct kinases that share similar consensus phosphorylation motifs (Reinhardt and Yaffe, 2009). Among others, Chk1, Chk2, and MK2 target cdc25 phosphatases, leading to their degradation or functional inactivation, which results in a rapid decrease of Cdk1 activity and suppression of cell cycle progression (Mailand et al., 2000; Peng et al., 1997; Reinhardt et al., 2007). In addition, Chk1/Chk2/MK2-independent pathways contribute to inhibition of mitotic kinases. For example, ATR-mediated degradation of the Plk1 co-factor Bora restricts Plk1 activity and ATM/ATR-mediated phosphorylation of p53 leads to the expression of the Cdk1 inhibitor p21 (Bunz et al., 1998; Qin et al., 2013).

Interestingly, not only does ATM/ATR inhibit mitotic kinases, but the mitotic kinases have also been implicated in reversing the checkpoint. Both Cyclin dependent kinase 1 (Cdk1) and Pololike kinase 1 (Plk1) phosphorylate multiple targets in the DDR (van Vugt et al., 2010). In particular Plk1-mediated degradation of Claspin, a protein required for ATR-dependent Chk1 activation, severely affects checkpoint maintenance (Mailand et al., 2006; Mamely et al., 2006; Peschiaroli et al., 2006). Plk1 activation is tightly linked to the central cell cycle engine, where it in a feedback loop also stimulates Cdk1 activity (Lindqvist et al., 2009b). However, whereas Plk1 is redundant for unperturbed mitotic entry, it becomes essential for recovery from a DNA-damage arrest (van Vugt et al., 2004), suggesting that Plk1-mediated down-regulation of the DDR is a key function to allow recovery from a cell cycle checkpoint. A critical question therefore is what controls when Plk1 activity can start to accumulate during a DDR.

Here we show that the DNA damage-induced spread of ATM activity across chromatin prevents Plk1 activation. The ATM activity is efficiently counteracted by the chromatin-bound phosphatase Wip1, leading to Plk1 re-activation despite the presence of active ATM at DNA break sites and active ATR. Based on a mathematical model, we suggest that the G2 checkpoint does not function by monitoring completion of repair but rather that the global ATM/Wip1 balance on chromatin controls the minimal duration of a checkpoint arrest.

## Results

We constructed a FRET-based sensor that can respond to ATM and ATR activity and targeted it to chromatin by fusion with Histone H2B (referred to as H2B-ATKAR, Figure 1–figure supplement 1A-C). After treatment with the radiomimetic drug Neocarzinostatin (NCS) H2B-ATKAR specifically detects ATM activity (Figure 1–figure supplement 1C-E). The H2B-ATKAR signal was induced by NCS addition in a dose-dependent manner and decreased over time (Figure 1A and Figure 1–figure supplement 1F). To follow both ATM and Plk1 activities throughout a DDR, we next compared the dynamics of H2B-ATKAR and a Plk1 reporter (Macurek et al., 2008) from addition of NCS to spontaneous checkpoint recovery in single U2OS and p53-depleted RPE cells. Interestingly, the decrease of H2B-ATKAR signal coincided with reactivation of Plk1 that controls recovery from a DNA damage-dependent checkpoint, indicating that ATM may control the duration of a DDR (Figure 1B, C).

**Figure 1.**
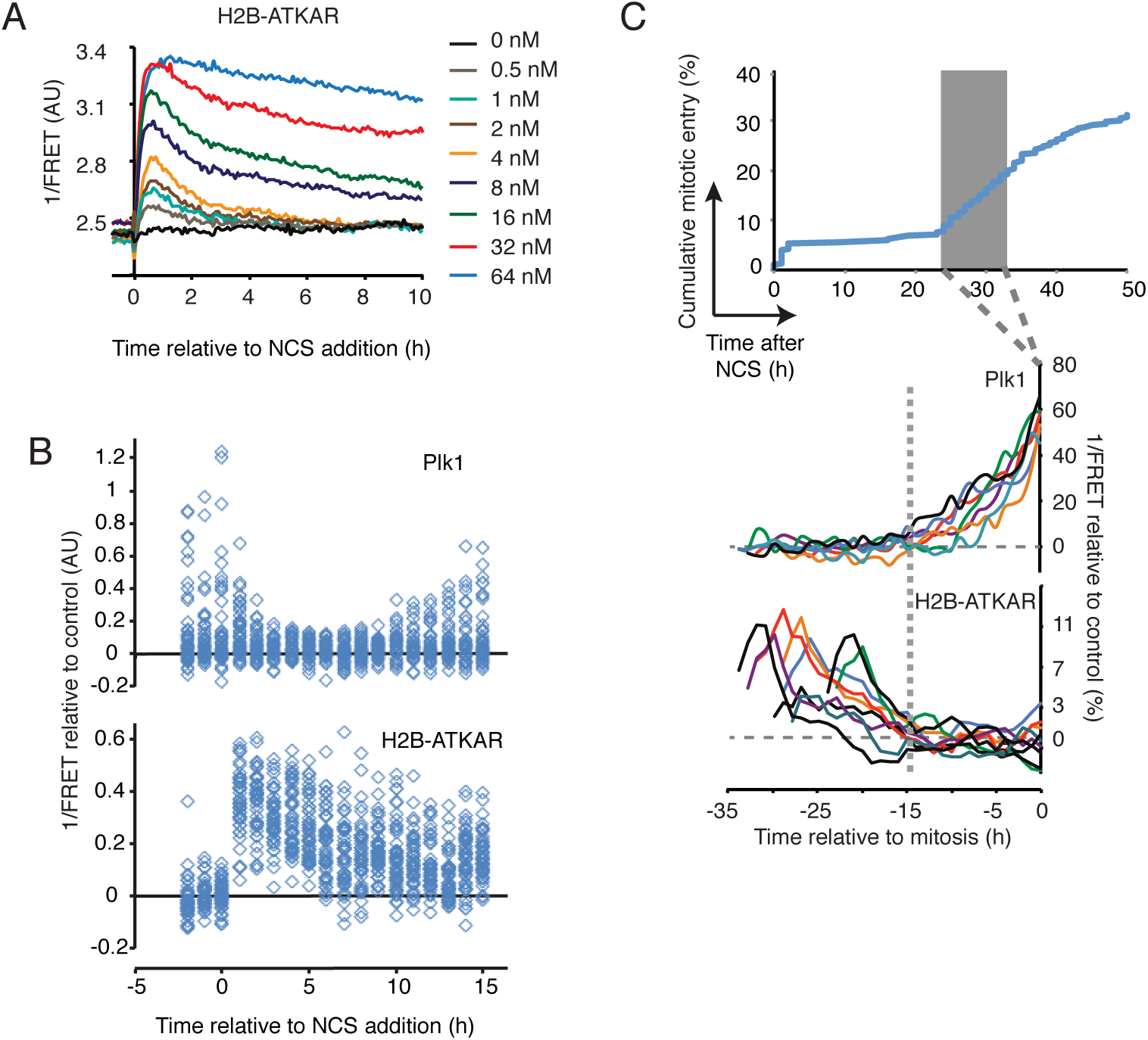
H2B-ATKAR dephosphorylation correlates to Plk1 activation during checkpoint recovery. (**A**) H2B-ATKAR signal responds in a dose-dependent manner to NCS addition and is reversed over time. Graph shows average 1/FRET of ≥10 U2OS cells/condition. (**B**) Resumption of Plk1 activity correlates with reversal of H2B-ATKAR phosphorylation. A mixed population of RPE cells expressing H2B-ATKAR or Plk1 FRET probe were transfected with p53 siRNA and treated with 8 nM NCS. 1/FRET was quantified of at least 41 cells per time point for each probe. H2B-ATKAR or Plk1 FRET were recognized by their nuclear or whole-cell localization. Each rectangle corresponds to one cell. **(C)** Reversal of H2B-ATKAR correlates with resumption of Plk1 activity during cell cycle restart. A mixed population of U2OS cells expressing H2B-ATKAR or Plk1 FRET probe were treated with 2 nM NCS and mitotic entry was followed over time (top). Cells entering mitosis 24 to 33h after NCS addition (gray rectangle) were synchronized *in silico* on mitosis and 1/FRET of individual cells was quantified (bottom).

To test if and when ATM controls Plk1 activation, we added a small molecule inhibitor to ATM at different time-points of a DDR. In accordance with a role for ATM to control Plk1 activity, inhibition of ATM in the early phases of a DDR sustained Plk1 activity (Figure 2A, B). In contrast, after Plk1 activity had restarted, its slow appearance (Liang et al., 2014) was counteracted by ATR (Figure 2C and Figure 2–figure supplement 1A). This suggests that both ATM and ATR control Plk1 activity, but that they function at different periods during a DDR. Indeed, ATM and ATR inhibition showed a synergistic effect on checkpoint duration only in the early phases of a DDR (Figure 2D, E and Figure 2–figure supplement 2B). Thus, our data is in support of a model where ATM controls when Plk1 activity can be initiated to switch off an ATR-dependent checkpoint (Figure 2F).

**Figure 2.**
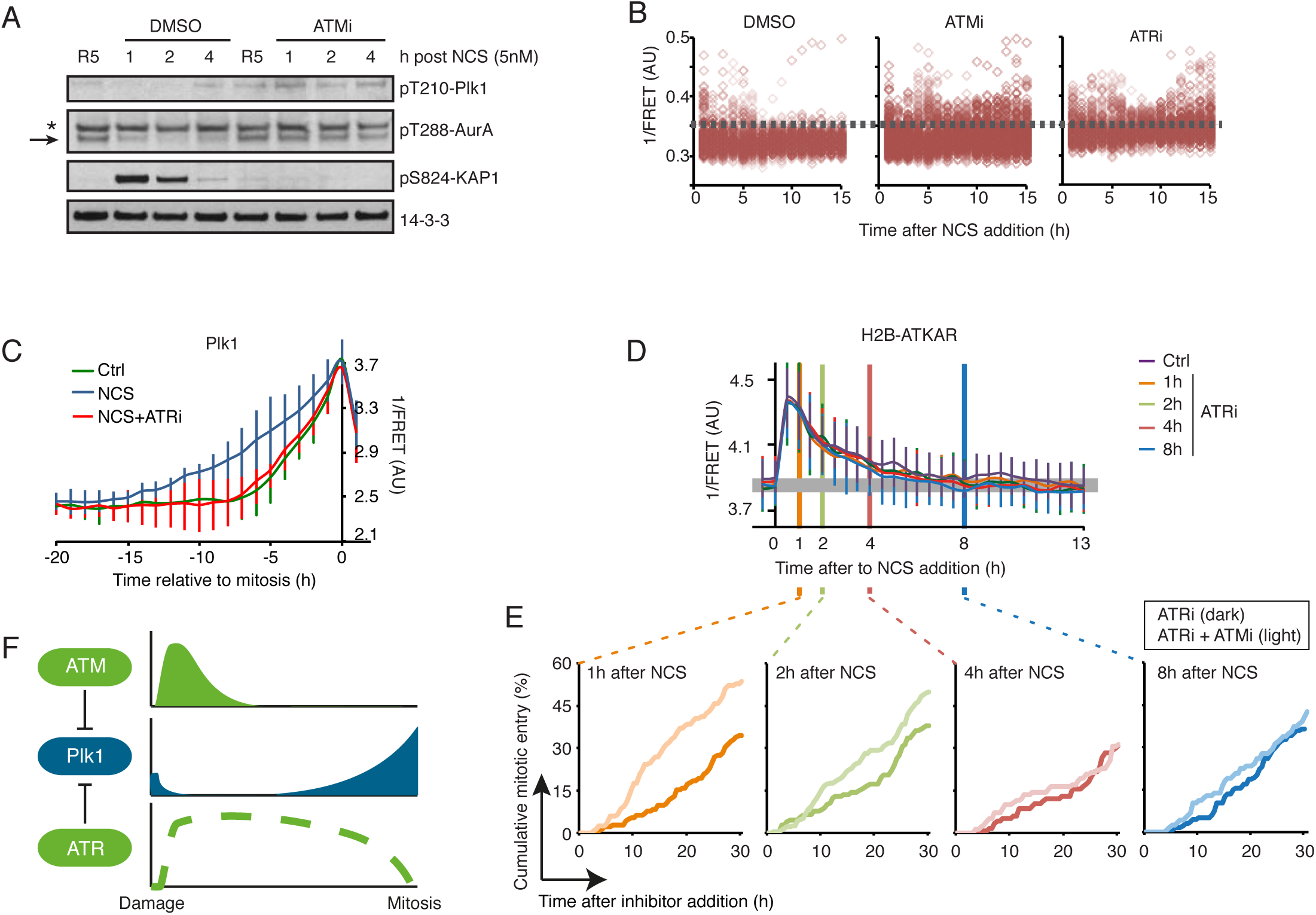
ATM inhibits Plk1 during the early phases of a DDR. (**A**) ATM inhibits Plk1 activity after NCS treatment. RPE cells were synchronized by 2 mM HU for 16 h and 5 h after release to fresh media treated with NCS (5 nM) and DMSO or ATMi (10 uM) for indicated times. Antibodies against pT210-Plk1 and pT288-Aurora A recognize active forms of Plk1 and Aurora-A, respectively. Asterisk indicates a cross-reacting band. (**B**) ATM activity contributes to Plk1 inhibition early after damage. U2OS cells expressing Plk1 FRET probe were treated with NCS (4 nM) and 15 min later ATMi, ATRi, or DMSO were added. Plots show 1/FRET of ~500 cells/condition/ time-point. (**C**) ATR counteracts Plk1 activity after cell-cycle re-start. Plk1 FRET-probe expressing U2OS cells were untreated (Ctrl) or treated with 2 nM NCS followed by DMSO or ATRi. 1/FRET of individual cells entering mitosis was quantified. Graph shows 1/FRET of ≥10 cells, synchronized *in silico* on mitosis. Note that the duration of Plk1 activation is longer in cells recovering from DNA damage compared to unperturbed cells, and that the prolonged duration is reverted in the presence of ATR inhibitor. (**D**) ATR inhibition does not affect H2B-ATKAR FRET-ratio after NCS. ATR inhibitor was added to U2OS cells expressing H2B-ATKAR at the indicated time points after 4 nM NCS addition. Graph shows average and SD of 1/FRET for ≥10 cells per condition. (**E**) Synergistic effect of ATM and ATR inhibition early after NCS. Cumulative mitotic entry of U2OS cells after treatment with NCS (1 nM) followed by addition of ATRi (1 uM, dark lines) or ATRi and ATMi (1 uM and 10 uM, light lines) at different time points as indicated. For comparison, these time-points are displayed as vertical lines in (D), where 4 nM NCS is used. (**F**) Schematic model. Whereas ATR inhibits Plk1 activity throughout a DDR, ATM determines when Plk1 can be activated to promote cell cycle resumption.

ATM is activated at sites of double strand breaks, and we next sought to assess whether H2B-ATKAR dephosphorylation and cell cycle re-start corresponds to ATM inactivation at DNA damage foci. We therefore established a setup where we followed the appearance of Plk1 activity in live cells and subsequently fixed and quantified immunofluorescence from the same cells (Figure 3–figure supplement 1). Interestingly, ATM-dependent phosphorylation of H2AX and p53 as well as autophosphorylation of ATM remained after Plk1 was re-activated, showing that ATM is active after initiation of cell cycle restart (Figure 3A-C). Moreover, repair proteins as 53BP1 and Rad51 were present in nuclear foci after Plk1 activation, indicating that H2B-ATKAR is dephosphorylated despite the presence of DNA breaks (Figure 3D). Sustained ATM activity after cell cycle re-start was also detected by ATKAR that is present in nucleoplasm, but is not targeted to chromatin. ATKAR shows faster and more sustained nuclear phosphorylation compared to H2B-ATKAR, suggesting a difference in phosphorylation dynamics of diffusible and chromatin-bound ATM targets (Figure 3-figure supplement 2). Importantly, ATKAR detects nuclear ATM activity throughout the recovery process, showing that ATM remains active until mitotic entry (Figure 3E-F). Thus, whereas ATM remains active throughout a DDR, H2B-ATKAR responds to a subset of ATM activity that is silenced before Plk1 re-activation.

**Figure 3.**
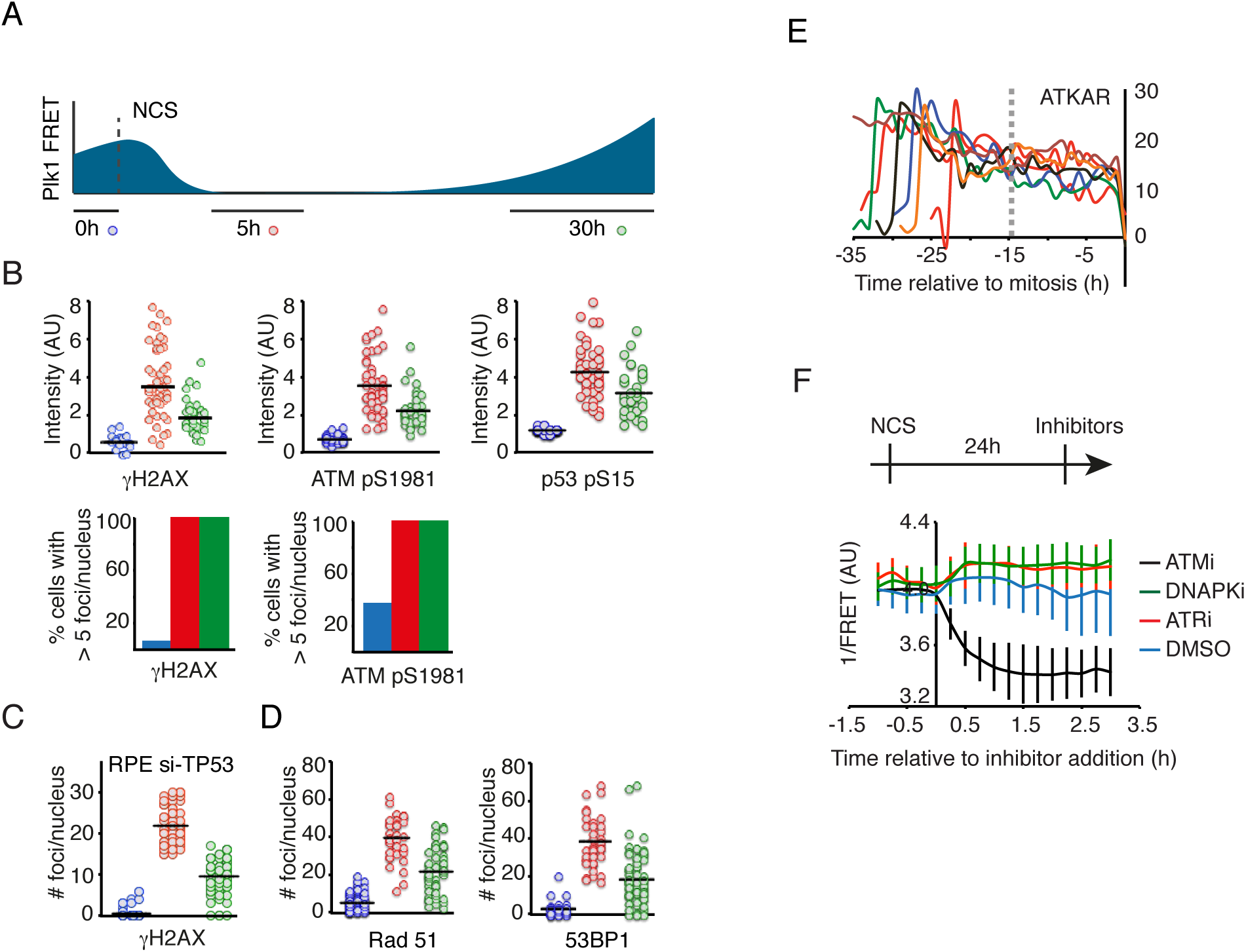
ATM is active after cell cycle re-start. (**A-D**) DNA damage foci are present after Plk1 re-activation. **(A)** Schematic of approach. U2OS cells expressing Plk1 FRET-probe were treated with NCS (2 nM). Before fixation at indicated time-points, 1/FRET was followed in individual live cells to detect undamaged G2 cells (0h, blue), G2 arrested cells without detectable Plk1 activity (5h, red), and recovering G2 cells with increasing Plk1 activity (30h, green). After fixation, the corresponding cells were identified both based on position and morphology. **(B)** Quantification of immunofluorescence of cells followed as in A. Graphs show signal intensity or percentage of cells with nuclear foci detected by indicated antibodies. Black bar indicates median and circles correspond to individual cells. **(C)** RPE cells expressing Plk1-FRET were treated with siRNA for TP53 and followed as in (A and B). Times were adjusted to 4.5 h (red) and 17 h (green). Graph shows quantification of γH2AX foci. **(D)** Quantification of immunofluorescence of cells followed as in A. Graphs show amount of nuclear foci detected by indicated antibodies. Black bar indicates median and circles correspond to individual cells. (**E**) ATKAR phosphorylation is sustained until mitosis during spontaneous checkpoint recovery. U2OS cells expressing ATKAR were followed during treatment with NCS (2 nM) and 1/FRET of cells spontaneously recovering 24 to 33 h later were plotted as in Figure 1C. Each line represents a single cell synchronized *in silico* upon mitotic entry. (**F**) ATKAR phosphorylation depends on ATM activity long after NCS addition. U2OS cells expressing ATKAR were treated with 5 nM NCS. The indicated inhibitors were added 24 h later. Graph shows average and SD of at least 15 cells.

As H2B-ATKAR is restricted to chromatin due to targeting by Histone H2B, this activity presumably occurs on chromatin. However, this activity is unlikely to be present on DNA damage foci, as we did not detect enrichment of H2B-ATKAR activity on DNA damage foci (not shown) and both yH2AX and pS1981 ATM staining persisted on foci after H2B-ATKAR signal disappeared (Figure 3B, C). Rather, we found that upon localized damage, H2B-ATKAR detected a global spread of ATM activity across chromatin (Figure 4A, B and Figure 4-figure supplement A, B). In contrast to the yH2AX and BRCA1 that remained restricted to close proximity of DNA lesions, H2B-ATKAR detected a chromatin-wide signal, indicating that ATM activity can reach chromatin far from the damaged area (Figure 4-figure supplement C, D). Taken together, this shows that ATM activity on DNA damage foci is not sufficient to block Plk1 re-activation, but rather suggests that spread of ATM activity across chromatin controls cell cycle restart.

**Figure 4.**
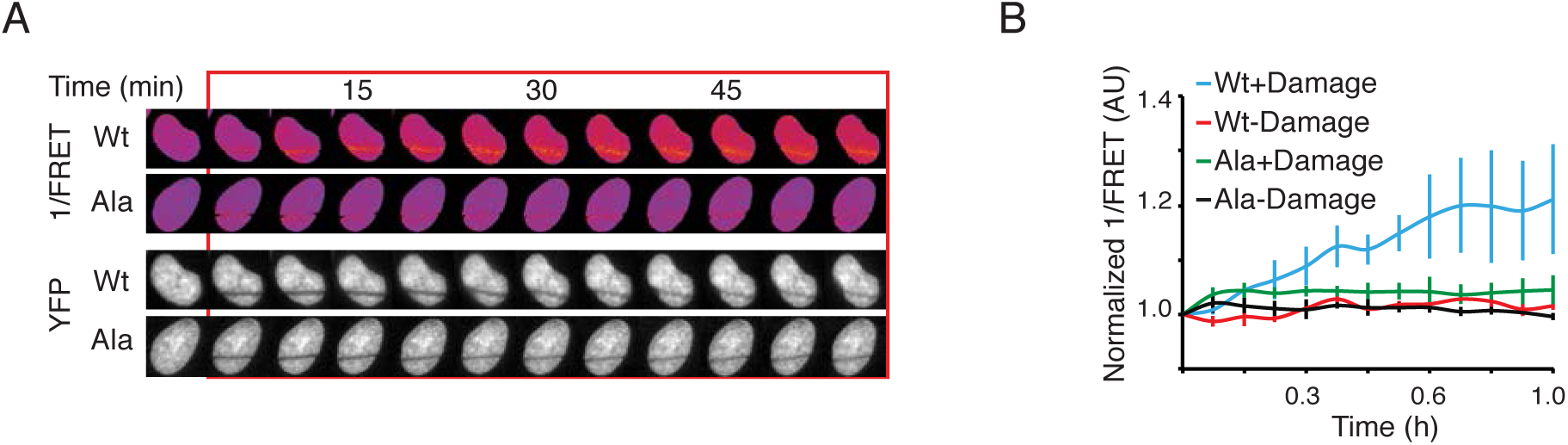
H2B-ATKAR detects spread of ATM activity over chromatin. **(A)** H2B-ATKAR 1/FRET spreads over chromatin after localized damage. U2OS cells expressing H2B-ATKAR or non-phosphorylatable Ala-H2B-ATKAR were laser-microirradiated and 1/FRET was followed over time. Note that bleaching of fluorophores precludes FRET-analysis within the laser-microirradiated area. **(B)** Quantification of spread of H2B-ATKAR FRET-change after laser microirradiation in U2OS cells. Measurements were performed distal to the laser-microirradiated area. Graph shows average and standard deviation of at least 7 cells per condition.

We next sought to identify why ATM-dependent phosphorylation of H2B-ATKAR on chromatin is more rapidly reverted compared to phosphorylation of diffusible ATKAR. We find that the spread of ATM activity across chromatin is controlled by the chromatin-bound phosphatase PPM1D (referred to as Wip1), which is known to counteract ATM-mediated phosphorylations (Macurek et al., 2010; Shreeram et al., 2006; Yamaguchi et al., 2007). Wip1 efficiently counteracted the H2B-ATKAR signal after NCS treatment, but less efficiently counteracted the diffusible ATKAR (Figure 5A, B). In addition, overexpression or depletion of Wip1 blocked or potentiated, respectively, the spreading of ATM activity at chromatin after localized DNA damage caused by laser microirradiation (Figure 5C, D). Whereas control cells were efficiently stimulated to enter mitosis once the H2B-ATKAR signal was reverted in the presence of an ATR inhibitor, Wip1-deficient cells did not revert the H2B-ATKAR signal nor enter mitosis (Figure 5E). This suggests that in addition to the established role of Wip1 to counter a p53-mediated cell cycle exit (Lindqvist et al., 2009a), Wip1 controls when ATM-mediated signaling throughout chromatin is reversed to allow initiation of a cell cycle re-start (Figure 5F).

**Figure 5.**
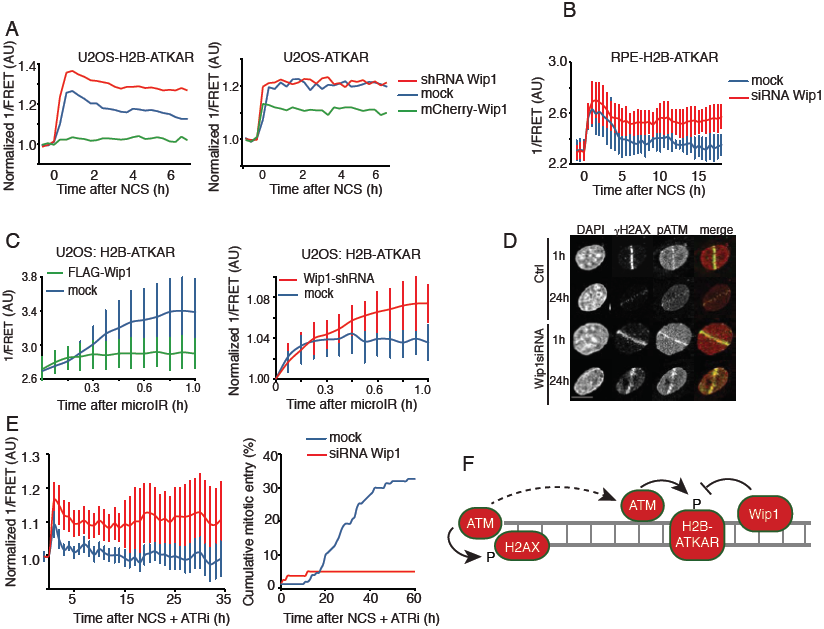
Wip1 counteracts ATM activity at chromatin. (**A**) Quantification of 1/FRET of mixed populations of U2OS cells expressing H2B-ATKAR or ATKAR. Cells were mock transfected or transfected with Wip1 shRNA or mCherry-Wip1 for 48 h and treated with NCS (8 nM). Graph shows average of ≥8 cells. (**B**) Quantification of 1/FRET of RPE-H2B-ATKAR transfected with control or Wip1 siRNA treated with NCS (8 nM). Graph shows average of≥8 cells, error bars indicate SD. (**C**) Wip1 influences the spread of H2B-ATKAR 1/FRET change. U2OS-H2B-ATKAR cells were transfected with mock (blue), FLAG-Wip1 (green) or Wip1 shRNA (red) and microirradiated with 364 nm laser. 1/FRET distal to the damaged area was quantified. Graph shows average and SD of at least 5 cells. (**D**) pS1981-ATM is present throughout chromatin and counteracted by Wip1. U2OS cells were transfected with control or Wip1 siRNA, fixed after 1 h or 24 h after microirradiation with 364 nm laser, and co-stained for γH2AX and pS1981-ATM. Scale bar: 15 um. (**E**) Wip1 depleted cells do not enter mitosis in presence of ATRi. 1/FRET (left) and cumulative mitotic entry (right) were measured in U2OS cells expressing H2B-ATKAR transfected with mock or Wip1 shRNA treated with NCS (4 nM) and ATRi. (**F**) Schematic model. Rather than DNA-damage foci, H2B-ATKAR signal detects ATM/Wip1 balance throughout chromatin.

To study possible implications of our findings, we assembled a mathematical model where ATM, Wip1, ATR, and Plk1-dependent pathways are treated as functional entities (Figure 6A). Simulating this model, we found that the balance of ATR and cell cycle activities determined the duration of a cell-cycle arrest, where rising self-amplifying cell cycle activities eventually forced inactivation of ATR. Importantly however, a spread of ATM activity on chromatin resets the initial cell cycle activities, thereby ensuring a delay before ATR-dependent activities could be inactivated (Figure 6B). The duration of this delay is likely not determined solely by ATR, as p53 and p38-dependent activities may influence the self-amplifying build-up of cell cycle regulators (Bunz et al., 1998; Reinhardt et al., 2010). Due to the spatial separation of ATM activation on DNA breaks and ATM function throughout chromatin, where Wip1 phosphatase efficiently dephosphorylates ATM targets, ATM activity throughout chromatin was rapidly reversed (Figure 6B). The speed of reversal depended on repair rate, and if damage was sustained at high levels a steady-state appeared with intermediate ATM-mediated phosphorylation, low cell-cycle activity and sustained ATR activation (Figure 6C). Testing the prediction on the influence of repair rate on ATM activity dynamics from the model, we note sustained H2B-ATKAR phosphorylation when interfering with DNA repair by PARP inhibition or RNF8 siRNA (Figure 6D). Importantly, below a threshold level of remaining damage, Plk1 activity could start to increase and eventually silence the checkpoint (Figure 4C). This is in line with our finding that repair factors are present at DNA damage foci after Plk1 activation (Figure 3A, B). We propose that spread of ATM activity on chromatin functions as a barrier that sets a timer. In this sense, the barrier not only defines a period when checkpoint recovery cannot occur, but by re-setting cell cycle activities it also ensures a delay before these activities can rise to override ATR-mediated signaling. The ATM-dependent barrier thereby ensures a minimal duration between its own silencing by Wip1 and mitotic entry. Thus, although a cell cannot efficiently sustain a G2 checkpoint in the presence of low amounts of damage, the timer ensures a considerable cell cycle delay during which DNA repair can occur.

**Figure 6.**
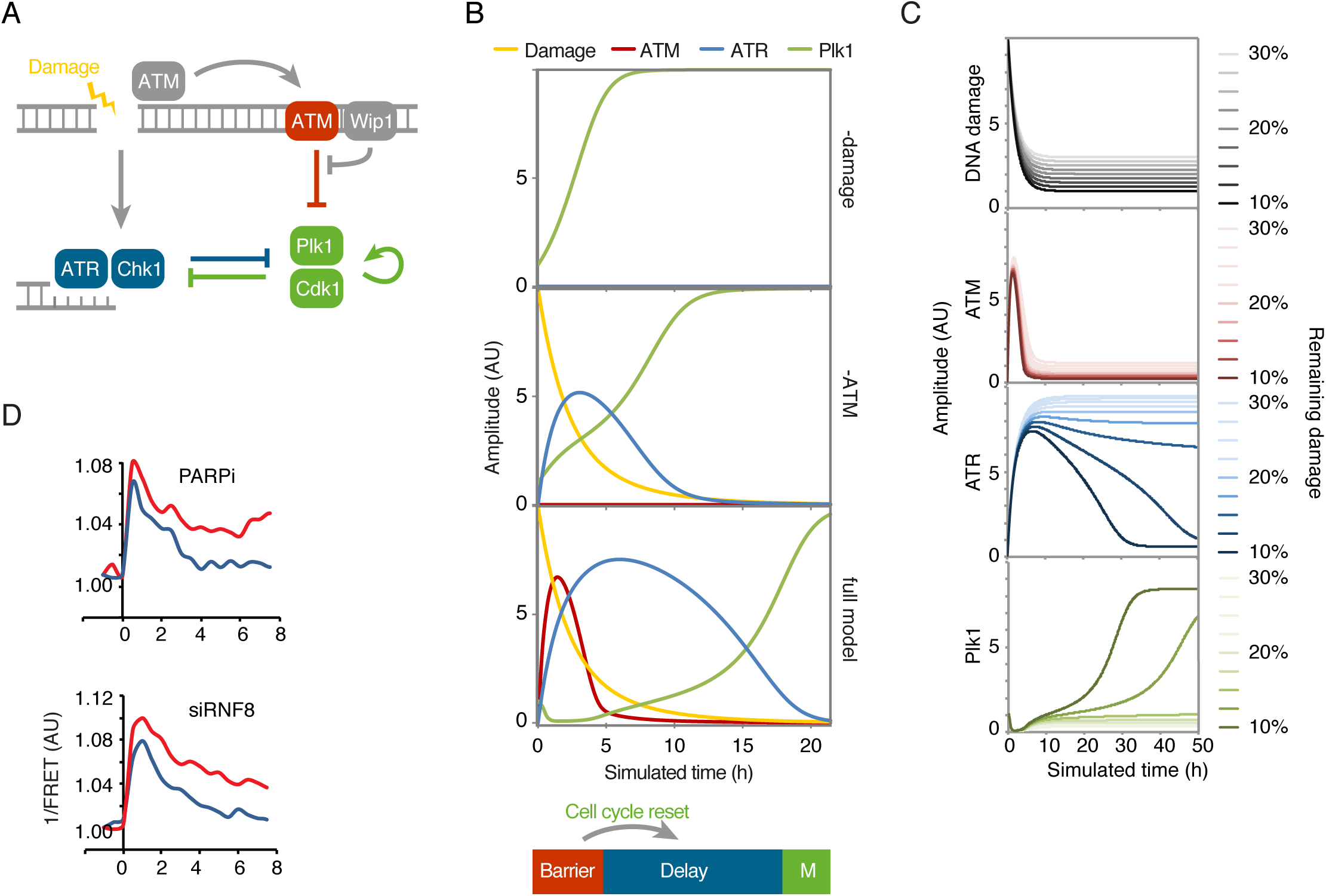
Wip1-dependent spread of ATM activity resets cell cycle signalling, thereby introducing a delay before ATR-dependent activities can be overcome. (**A**) Schematic outline of mathematical model. Arrows represent differential equations. (**B**) Simulation of model in the absence of damage (top) or ATM (middle) or containing all components (bottom). Spread of ATM activity functions as a barrier that blocks Plk1 activity. Wip1 efficiently counteracts ATM-mediated phosphorylations, which restricts spread of ATM activity to high damage levels. After reversal of the barrier, cell cycle signalling eventually overcomes ATR-dependent activities, despite the presence of unrepaired DNA breaks. Due to the reset cell cycle activities, a delay is introduced before ATR activities are overcome and mitosis occurs. (**C**) Cell cycle progression depends on a threshold level of damaged DNA. Simulation of model depicted in A, but set so that 10 - 30% of initial damage is not repaired, as shown in top graph. Above a threshold level of damage, ATM activity remains sufficiently high to together with ATR ensure that a cell cycle restart will not occur. (**D)** Interference with DNA repair processes delays dephosphorylation of H2B-ATKAR. 1/FRET was quantified in U2OS cells expressing H2B-ATKAR treated with 4 nM NCS in the presence of PARP inhibitor or RNF8 siRNA. Graphs show average of at least 10 cells.

We next sought to assess through which endogenous substrates ATM enforces a barrier to cell cycle restart. We found that similar to H2B-ATKAR, DNA damage-induced phosphorylation of Kap1-S824 and SMC3-S1083 spread throughout chromatin in an ATM-dependent manner and rapidly declined due to dephosphorylation by Wip1 (Figure 7A, B, and Figure 7–figure supplement A, B). This is in accordance with previously reported chromatin-wide effects of Kap1 and SMC3 after DNA damage (Kim et al., 2010; Ziv et al., 2006). Focusing on Kap1, we found that Wip1 interacted with Kap1 and efficiently dephosphorylated Kap1 S824 *in vitro* and *in situ* (Figure 7C, D, and Figure 7–figure supplement C). Moreover, inhibition of ATM in the absence of Wip1 sustained Kap1 S824 phosphorylation, indicating that Wip1 does not cause Kap1 dephosphorylation by impeding ATM function (Figure 7–figure supplement D). In addition to Wip1, we found that PP4 to some extent contributed to Kap1 dephosphorylation in untransformed RPE cells (Figure 7–figure supplement E, F) (Bulavin et al., 2002; Lee et al., 2012; Rauta et al., 2006). In contrast to ATM targets as H2AX, Kap1 S824 was dephosphorylated before cell cycle resumption was initiated (Figure 3B, 7E and 7F). Thus, phosphorylation of Kap1 S824 and H2B-ATKAR both depend on ATM and Wip1 and follow similar spatiotemporal dynamics. Importantly, Kap1-S824A expression stimulated cell cycle restart after DNA-damage, showing that phosphorylation of Kap1 can influence the duration of a DDR (Figure 7G). Taken together, ATM and Wip1-dependent regulation of Kap1 contributes to determine the duration of a cell cycle arrest, most likely in concert with other ATM and Wip1 targets.

**Figure 7.**
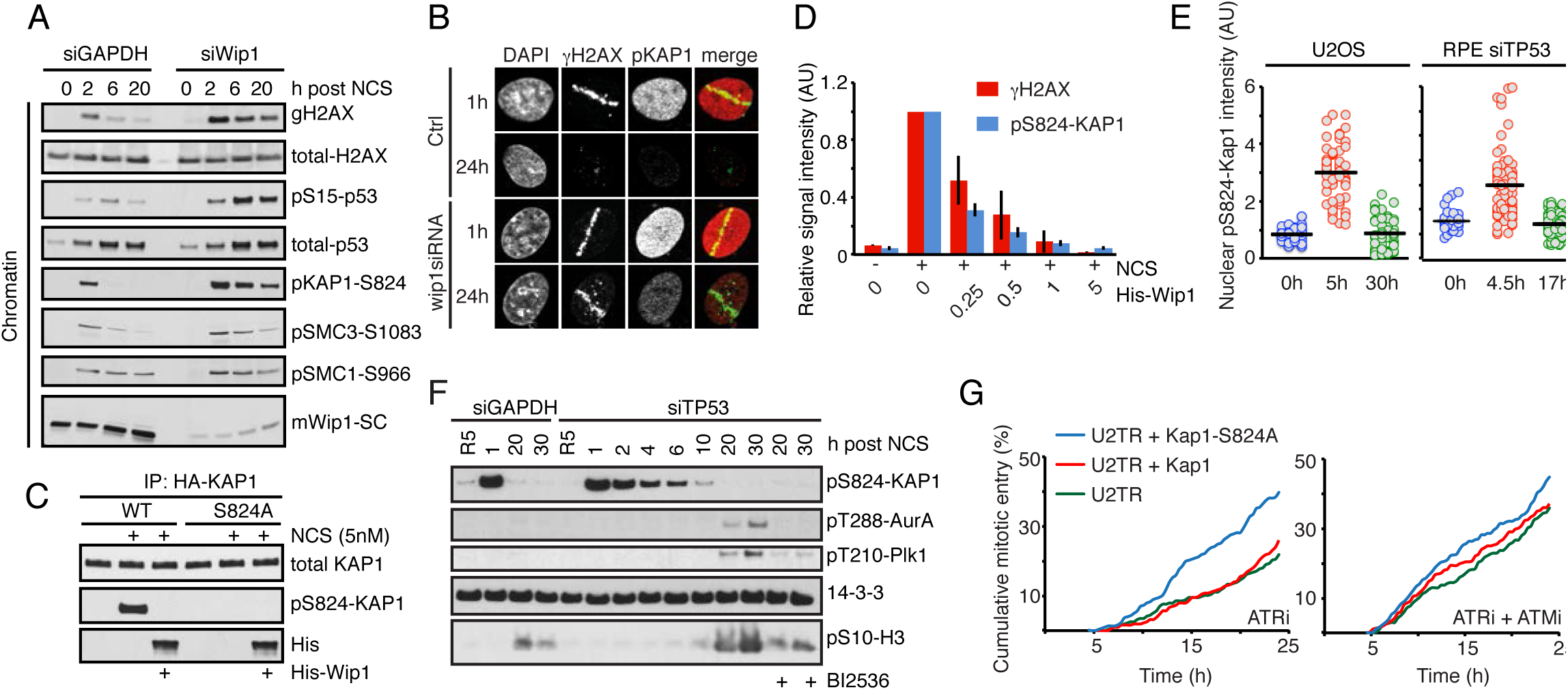
Kap1 is an ATM/Wip1 target on chromatin. (**A**) U2OS cells transfected with GAPDH or Wip1 siRNA were treated with NCS (5 nM) and collected after 2, 6 and 20 h. Chromatin fractions were probed with indicated antibodies. (**B**) RPE cells transfected with Wip1 siRNA were microirradiated, fixed 1 or 24 h later, and stained with the indicated antibodies. (**C)** HA-KAP1-WT or-S824A were immunopurified from cells exposed to NCS, incubated with His-Wip1 and probed with pS824-KAP1 or KAP1 antibody. **(D)** U2OS cells were fixed 1 h after treatment with NCS, incubated with His-Wip1 (0-5ng/ul) and probed for gH2AX and pS824-KAP1. Plot shows mean nuclear fluorescence intensity, error bars indicate SD. **(E**) Kap1 is dephosphorylated before Plk1 activation. RPE cells transfected with TP53 siRNA and U2OS cells were followed as in Fig. 2A and stained for pS824-Kap1. For RPE cells, the times were modified as indicated. **(F)** RPE cells transfected with GAPDH or TP53 siRNA were synchronized by HU, released to fresh media for 5 h (R5) and treated with NCS for indicated times. Nocodazole (NZ) was added 1 h after NCS. Where indicated, cells were incubated in the presence of BI2536. Whole cell lysates were probed with indicated antibodies. (**G**) Overexpression of Kap1-S824A phenocopies ATM inhibition. Cumulative mitotic entry of ≥300 U2OS cells expressing inducible HA-tagged Kap1-wt (red) or Kap1-S824A (green) after treatment with NCS (4 nM) and subsequent treatment after 1h with ATRi or ATRi + ATMi.

## Discussion

Our results suggest that the duration of a cell cycle arrest is determined by three principal components. First, an ATM-dependent signal that efficiently blocks cell cycle progression. This signal is not affected by cell cycle regulators, but is rapidly reversed by the phosphatase Wip1. Second, an ATR dependent signal that counters cell cycle progression throughout the cell-cycle arrest, and third, the self-amplifying mitotic entry network that counters the ATR-mediated pathway in G2 phase.

Plk1 activity rises through G2 phase due to the self-amplifying properties of the mitotic entry network (Akopyan et al., 2014; Lindqvist et al., 2009b). Maintaining a self-amplifying activity at a constant level is not a trivial task for a cell. Indeed, we recently found that during a DDR, Cyclin B1 levels over time accumulate to levels far higher than is observed during an unperturbed cell cycle (Mullers et al., 2014). Thus, although ATR-mediated activities slow down cell cycle progression, G2 activities may ultimately prevail and induce mitotic entry. In particular Plk1 phosphorylates a large number of proteins involved in both DDR and cell cycle control and is essential for recovery from a cell cycle arrest in G2. By blocking Plk1 activity, ATM thereby not only solidly enforces a cell cycle block, but also ensures that mitotic entry will be postponed to long after the ATM-dependent signal is reversed.

The spatial regulation of the DDR on chromatin has recently attracted considerable attention (Altmeyer and Lukas, 2013). ATM activity is efficiently induced at damage sites, where it triggers establishment of DNA damage-induced foci. A determinant of foci formation is ATM-mediated phosphorylation of H2AX, which remains restricted to the damaged area, at least in part by insulation mediated by Brd4 (Floyd et al., 2013). Although DNA damage foci are likely to be crucial for repair and amplification of the DDR, the localized DNA damage needs to be communicated to the cell cycle machinery at a cellular scale. Here, we show that ATM activity is differentially maintained at subnuclear compartments, where ATM activity present at chromatin distal to damage sites blocks cell cycle progression. Whereas Wip1 may not be sufficient to abolish ATM activity in DNA-damage foci until damage is repaired, Wip1 efficiently counters the spread of active ATM to undamaged regions of chromatin. In this sense, ATMs stimulatory effect on repair may be sustained while ATMs effect on the cell cycle is reversed. We suggest that separating activating and inactivating locations may be a powerful manner for a cell to simultaneously construct a timer and a sustained signal.

In unicellular organisms, a checkpoint can eventually be overcome, despite that not all DNA lesions are repaired. In budding yeast, this process called adaptation depends heavily on the Plk1 homolog cdc5, which counteracts activating phosphorylation of the checkpoint kinase Rad53 (Donnianni et al., 2010; Toczyski et al., 1997). In contrast, multicellular organisms have evolved mechanisms that promote apoptosis or terminal cell cycle arrest, largely dependent on p53 (Bartek and Lukas, 2007; Belyi et al., 2010). We find that albeit the kinetics may differ, ATM-Wip1-Kap1 proteins function similarly in U2OS and RPE cells. However, untransformed G2 cells are more likely to terminally exit the cell cycle, rather than to recover from DNA damage. Strikingly, the duration of the ATM activity present across chromatin overlaps with the time required for a DNA damage-mediated cell cycle exit to become irreversible (Krenning et al., 2014; Mullers et al., 2014). We suggest that a timer is inherent to all cells, but that its effect will only become apparent upon deregulation of the p53 pathway. Of note, our model also elucidates the previously unexplained observations that cells can enter mitosis in the presence of low levels of damaged DNA (Lobrich and Jeggo, 2007). Such revival of adaptation may have deleterious consequences in multicellular organisms since segregation of unrepaired DNA during mitosis may cause aneuploidy and cancer.

## Methods

### Construction of ATM/ATR kinase activity reporter (ATKAR)

ATKAR is based on an established Plk1 biosensor (Fuller et al., 2008; Macurek et al., 2008), where the Plk1 target sequence from the parental biosensor was replaced with EPPLTQEI sequence derived from the residues 11-18 of human p53, a well established substrate of ATM and ATR kinases. To promote binding of the target site by FHA2 domain in the biosensor, the serine corresponding to Ser15 in p53 sequence was replaced by threonine and residue at +3 position was changed to isoleucine (Durocher et al., 2000). The DNA fragment corresponding to ATKAR was inserted in HindIII/XbaI sites of pcDNA4 plasmid. To generate H2B-ATKAR, DNA fragment carrying H2B sequence was cloned in-frame with ATKAR into the HindIII site. In contrast to a previously described ATM biosensor (Johnson et al., 2007), the FRET-ratio change observed after NCS addition was largely reversed upon ATM inhibition, indicating that phosphorylation of ATKAR is reversible and depends on ATM activity also after initiation of a DDR.

### Plasmids

Plasmid carrying human HA-KAP1 was obtained from Addgene (ID: 45569, (Liang et al., 2011)). The S824A mutant of HA-KAP1 was generated by Site-directed mutagenesis kit (Agilent Technologies) and mutation was confirmed by sequencing. Fragments corresponding to HA-KAP1-WT and HA-KAP1-S824A were cloned into pcDNA4/TO plasmid. pRS-Wip1 plasmid for shRNA-mediated knock-down of Wip1 was described previously (Lindqvist et al., 2009a).

### Cell culture and transfections

U2OS, MCF-7 and HEK293 cell lines were cultured in Dulbecco’s modified Eagle’s medium (DMEM) + GlutaMAX (Life Technologies) supplemented with 6% or 10% heat-inactivated fetal bovine serum, respectively (FBS, Hyclone) and 1% Penicillin/Streptomycin (Hyclone) at 37°C and 5% CO_2_. hTERT-RPE1 cell lines were cultured in DMEM/Nutrient mixture-F12 medium (DMEM/F12) + GlutaMAX (Life Technologies) supplemented with 10% heat-inactivated fetal bovine serum and 1% Penicillin/Streptomycin (Hyclone) at 37°C and 5% CO_2_. To generate cells stably expressing ATKAR or H2B-ATKAR, cells were transfected by linearized plasmids and selected by Zeocine for 3 weeks. For RNA interference experiments cells were seeded at a density of 8000 cells/well and transfected with SMARTpool ON-TARGET plus siRNAs (20 nM, Dharmacon) targeting Wip1, ATR, and p53 using HiPerFect (Qiagen) and OptiMEM (Life Technologies) at 48 h before analysis of phenotypes. Alternatively, cells were transfected with Silencer Select siRNA (5 nM, Life Technologies) targeting GAPDH (control), Wip1 (CGAAAUGGCUUAAGUCGAA), TP53 (GUAAUCUACUGGGACGGAA) or PP4C (UCAAGGCCCUGUGCGCUAA) using RNAiMAX (Life Technologies) and cells were analyzed 48 h after transfection, U2OS cells expressing KAP1 or KAP1-S824A upon tetracycline induction were generated as described previously (Macurek et al., 2008). Where indicated, hRPE cells were synchronized in S phase by hydroxyurea (2 mM, 16 h), released to fresh media for 5 h to allow progression to G2 and treated with NCS for indicated times.

### Live-cell microscopy and Image processing

For live cell imaging 8000-10000 cells were seeded in 96-well imaging plates (BD Falcon) 24 h before imaging in CO2-independent medium (Leibovitz’s L15- Invitrogen) supplemented with 6% or 10% heat-inactivated fetal bovine serum. Cells were followed on either a DeltaVision Spectris imaging system (Applied Precision) using a 20X, NA 0.7 objective, a Leica DMI6000 Imaging System using a 20X, NA 0.4 objective, or on an ImageXpress system (Molecular Devices) using a 20X, NA 0.45 objective. Images were processed and analyzed using ImageJ (http://rsb.info.nih.gov/ij/) or using custom written Matlab scripts. 1/FRET was quantified as the ratio of YFP emission-YFP excitation and CFP excitation-YFP emission as described previously (Hukasova et al., 2012). Unless stated otherwise, the median pixel value of the inversed nuclear FRET-ratio was used. For spontaneous recovery of H2B-ATKAR expressing cells, a moving average of 3 or 4 time-points is shown.

### Microirradiation

U2OS or RPE cells, grown on MatTek glass bottom dish (MatTek Corporation) were treated with 10 **u**M BrdU (Sigma) for 24 h. For micro-irradiation, the dish was mounted on stage of Leica DMI 6000B microscope stand (Leica) integrated with a pulsed nitrogen laser (20 Hz, 364 nm, Micropoint Ablation Laser System) that was directly coupled to the epifluorescence path of the microscope and focused through a Leica 40X HCX PL APO/1.25-0.75 oil-immersion objective. Typically, 50 cells were micro-irradiated (150X 1pixel) in Leibovitz’s L15 medium at 37 °C, after which cells were either followed for 1 h or returned to incubator at 37°C to recover. Cells were fixed 1 h or 24 h after microirradiation. Fixed samples were analyzed on Zeiss LSM510 META confocal microscope equipped with a 63X Plan-A (1.4 NA) oil-immersion objective. Images were recorded using Zeiss LSM imaging software in multi-track mode.

### Immunofluorescence

For immunofluorescence, U2OS or RPE cells were seeded on 96-well microscope plates (Falcon-BD), or MatTek glass bottom dishes. Fixation was performed using 3.7% paraformaldehyde (Sigma) for 5 min at room temperature. Permeabilization was achieved by incubating cells with ice-cold methanol for 2 min. Blocking, antibody and DAPI incubations were performed in TBS supplemented with 0.1% Tween-20 (TBST) and 2% BSA (Sigma). Wash steps were performed in TBS supplemented with 0.1% Tween-20. Images were acquired using either DeltaVision Spectris imaging system using a 20X, NA 0.7 objective, a Leica DMI6000 Imaging System using a 20X, NA 0.4 objective, or on an ImageXpress system using a 20X, NA 0.45 objective and quantified as described (Akopyan et al., 2014).

### Immunoprecipitation

U2OS cells expressing ATKAR or H2B-ATKAR were collected 1 h after exposure to IR (5 Gy) or UVC (10 J/m2) or NCS (5 nM) and sonicated in cold IP buffer (50 mM HEPES pH 7.5, 250 mM NaCl, 0.25 NP-40, 1% glycerol) supplemented with phosphatase inhibitor (PhosSTOP, Roche) and protease inhibitor (complete EDTA-free, Roche). FRET probes were immunoprecipitated from cell extracts using GFP-Trap (ChromoTek) for 2 h. Beads were washed 3 times with IP buffer and mixed with SDS sample buffer. Alternatively, HEK293 cells were transfected by empty EGFP or EGFP-Wip1 plasmid using linear polyethylenimine MAX. After 48 h cells were harvested to lysis buffer (50mM Tris pH 7.5, 150mM NaCl, 3 mM MgCl2, 10% glycerol, 1% Tween-20, 0.1% NP-40) supplemented with benzonase (25 U/ml), protease and phosphatase inhibitors (Roche), sonicated and incubated on rotary shaker overnight at 4°C. Cell extract was centrifuged 15 min 20000g at 4°C, supernatant was incubated with GFP-Trap beads for 1 h at 4°C, beads were washed four times with lysis buffer and bound proteins were eluted with SDS sample buffer.

### Subcellular fractionation

Cells were treated with DMSO or 5 nM NCS and collected after 2, 6 or 20 h. Subcellular fractionation was performed as described before (Macurek et al., 2010). Cells were incubated in buffer A (10 mM HEPES, pH 7.9, 10 mM KCl, 1.5 mM MgCl2, 0.34 M sucrose, 10 % glycerol, 1 mM DTT, 0.1% Triton X-100 and protease inhibitor cocktail) at 4°C for 10 min and centrifuged at 1500 g for 2 min. The cytosolic fraction was collected and sedimented cells were further incubated with buffer B [10mM HEPES, pH 7.9, 3 mM EDTA, 0.2 mM EGTA, 1 mM DTT] and centrifuged at 2000 g for 2 min. Nuclear soluble fraction was collected, pooled with cytosol (together forming a soluble fraction) and mixed with 4x SDS sample buffer. Remaining chromatin fraction was mixed with 1.25 x volumes of 1x SDS sample buffer. All samples were boiled, sonicated and separated on SDS-PAGE.

### In vitro and in situ phosphatase assay

Cells expressing HA-KAP1-WT or HA-KAP1-S824A were treated with DMSO or NCS (5 nM) for 1 h and then extracted using IP buffer. Cell extracts were immunoprecipitated for 2 h using monoclonal anti-HA tag antibody (Santa Cruz, sc7392) immobilized at pA/G beads (Pierce). Beads were stripped with 0.5 M NaCl to remove proteins interacting with KAP1. Beads were incubated with purified His-Wip1 (100 ng) in phosphatase buffer (40 mM HEPES pH 7.4, 100 mM NaCl, 50 mM KCl, 1 mM EGTA, 50 mM MgCl2) for 20 min at 30 °C. The reaction was stopped by addition of 4x SDS sample buffer. Alternatively, cells grown on coverslips were treated with NCS (5 nM) for 1 h, fixed with 4 % paraformaldehyde, permeabilized with 0.5 % Triton X-100 and *in situ* phosphatase assay was performed as described (Munoz et al., 2013). Samples were blocked with 3 % BSA in PBS and incubated at room temperature in ISB buffer (50 mM HEPES pH 7.0, 10 mM MgCl2, 25 mM NaCl, 2 mM CaCl2, 1 mM DTT) and His-Wip1 (0.25 - 10 ng/ul) for 1 h. Reaction was stopped by addition of 20 mM NaF and 20 mM β-glycerolphosphate in PBS. Samples were incubated with a mixture of anti-gH2AX and anti-pS824-KAP1 antibodies for 1 h. After washing samples were incubated with secondary antibodies and DAPI. Imaging was performed on ScanR microscope (Olympus) and 1000 nuclei were counted per condition.

### Reagents and antibodies

The following antibodies were from Cell Signaling: pHistone H3 (#3377); pChk1-S345 (133D3; #2348); pATM-S1981 (10H11.E12; #4526); pH2AX (20E3; #9718), pp53-s15 (#9284; #9286), Phospho-(Ser/Thr) ATM/ATR Substrate (#2851), pT210-Plk1 (#9062), pT288-AurA (#3079), AurA (#3092), pS296-Chk1 (#2349), pS317-Chk1 (#12302). Antibodies against pSMC3-S1083 (#IHC00070), pKap1-S824 (#A300-767A and GTX63711), Kap1 (#A300-274A and GTX62973), pS966-SMC1 (A300-050A), and PP4C (GTX114659) were from Bethyl Lab and GeneTex. Additional antibodies included pH2AX (Clone JBW301; #05-636, Millipore), 53BP1 (#NB100-304, Novus Biologicals.), pS10-H3 (#05-806, Millipore), Rad51 (FE#7946, Biogenes), and Alexa-and FITC-coupled antibodies (Life Technologies). Antibodies against GFP (sc-8334), 14-3-3, p53 (sc-6243), p21 (sc-397), Wip1 (sc-130655), His (sc-8036) and BRCA1 (sc-6945) were obtained from Santa Cruz. Neocarzinostatin (NCS), Etoposide and DMSO were from Sigma. ATM inhibitor (#4176 and #3544) and DNA-PK inhibitor (#2828) were from Tocris Bioscience and used at 10 uM. ATR inhibitor VE821 (Reaper et al., 2011) was obtained from (Tinib-Tools) or was synthesized according to published procedure (Charrier et al., 2011) and used at 1 uM. PARP inhibitor KU0058948 was used at 10 uM. Plk1 inhibitor BI2536 was from Boehringer Ingelheim Pharma and used at 100 nM. CDT was a generous gift from Teresa Frisan.

### Simulations

We wrote ordinary differential equations assuming Michaelis-Menten kinetics to simulate the relation between the variables Damage, Cell Cycle, ATM, and ATR. Damage was added at the beginning of a simulation and assumed to decrease over time, simulating quick repair by non-homologous end joining and slow repair by homologous recombination. Cell Cycle contains a positive feedback loop to simulate the auto-amplifying property of the mitotic entry network, of which Plk1 is a part. ATM is activated by damage, but is efficiently counteracted by a constant to simulate the balance of ATM and Wip1 on chromatin. ATR, simulating ATR-Chk1 activity, is activated by Damage, but inactivated by Cell Cycle to simulate Plk1 and Cdk1 phosphorylation of DDR components. Both ATM and ATR inhibit Cell Cycle.

The equations were solved using Copasi 4.8, build 35 (www.copasi.org/). Parameterization was performed manually, restricting constants to 1, 2, or 10.

### Differential equations

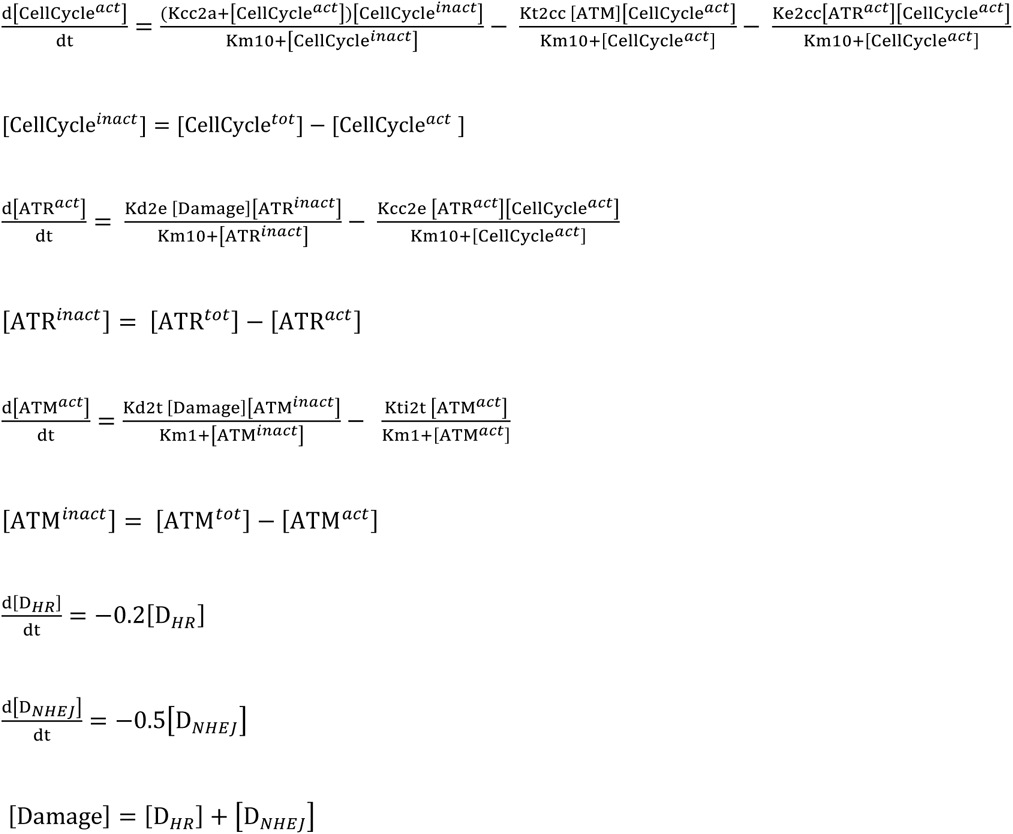

### Parameters

| Name | Type | Value |
| --- | --- | --- |
| Kcc2a | fixed | 1 |
| Kcc2e | fixed | 1 |
| Kd2e | fixed | 1 |
| Kd2t | fixed | 2 |
| Ke2cc | fixed | 1 |
| Km1 | fixed | 1 |
| Km10 | fixed | 10 |
| Kt2cc | fixed | 10 |
| Kti2t | fixed | 10 |

| Name | Type | Initial Value |
| --- | --- | --- |
| CellCycle<sup>act</sup> | ode | 1 (10% active) |
| CellCycle<sup>tot</sup> | ode | 10 |
| D<sub>HR</sub> | ode | 3 (30% HR) |
| D<sub>NHEJ</sub> | ode | 7 (70% NHEJ) |
| ATR<sup>act</sup> | ode | 0 |
| ATR<sup>tot</sup> | ode | 10 |
| ATM<sup>act</sup> | ode | 0 |
| ATM<sup>tot</sup> | ode | 10 |

## Acknowledgments

We thank Marcel van Vugt, Camilla Sjögren, and Urban Lendahl for comments on the manuscript. This work was supported by grants from the Swedish Research Council, the Swedish Foundation for Strategic Research, the Swedish Cancer Society and the Swedish Childhood Cancer Foundation (all to AL), Vinnova (TH), the Grant Agency of the Czech Republic (13-18392S) and Ministry of Education, Youth and Sports (PHOSCAN). HJ was supported by the Wenner-Gren foundation and JB partly by the Grant Agency of the Charles University (project 836613).

---

**Figure 1–figure supplement.**
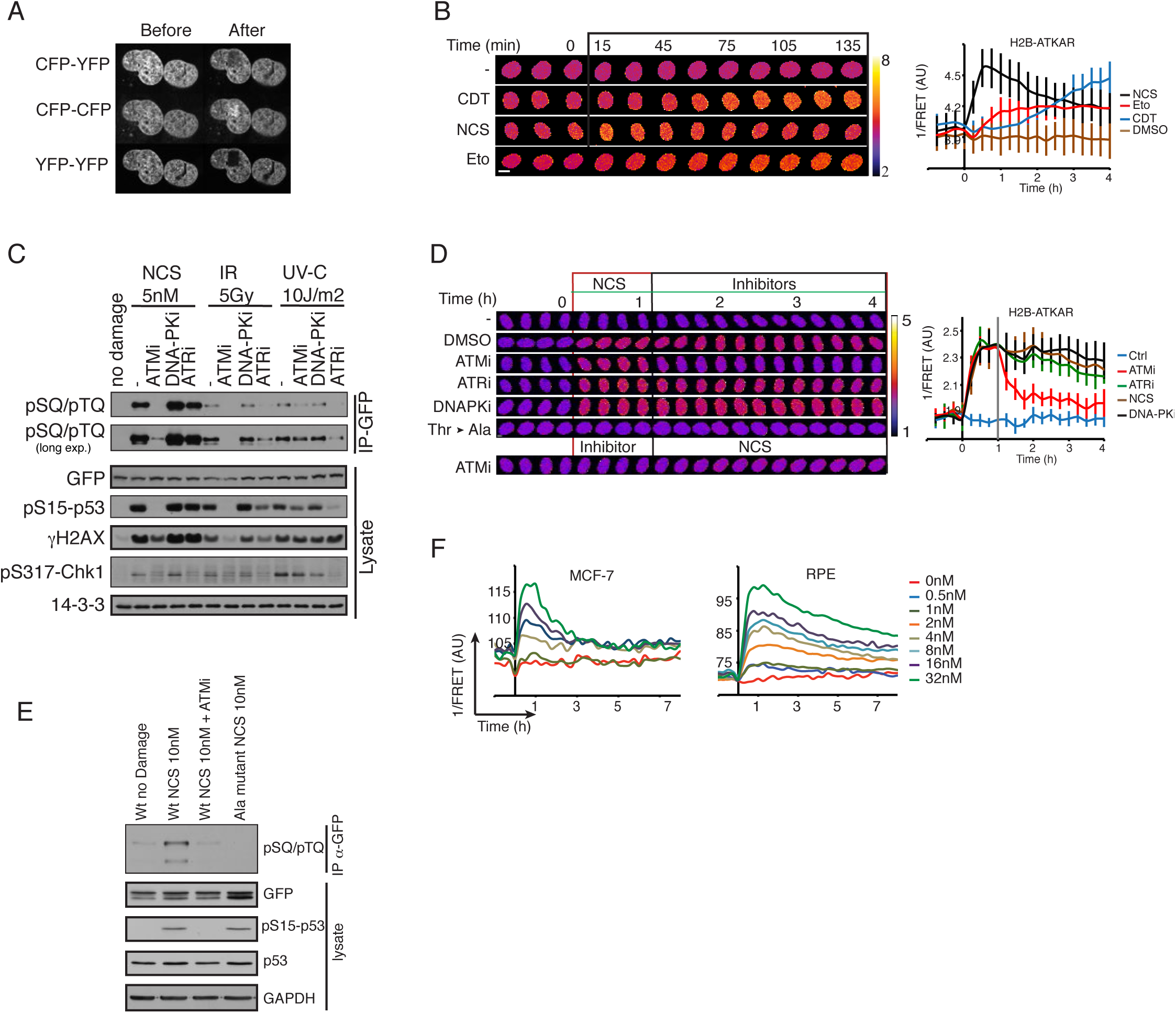
A FRET based biosensor to monitor the activity of ATM/ATR kinase in live cells. (**A**) Acceptor photobleaching of H2B-ATKAR. U2OS cells expressing H2B-ATKAR were photobleached using a 514 nm laser and images were acquired by using CFP-YFP, YFP-YFP and CFP-CFP excitation-emission before and after photobleaching. The bleached area is visible in the YFP-YFP images. (**B**) Kinetics of H2B-ATKAR 1/FRET change after treatment with Etoposide (Eto), Neocarzinostatin (NCS) or cytolethal distending toxin (CDT). Time-lapse sequence (left) or quantification of 1/FRET (right) of U2OS cells expressing H2B-ATKAR. Graph shows average and SD of at least 15 cells. Time point 0 indicates addition of drugs. (**C**) H2B-ATKAR phosphorylation after NCS addition depends on ATM. GFP pull-down from U2OS cells expressing H2B-ATKAR treated with NCS (5 nM) or exposed to IR (5 Gy) or UVC (10 J/m2). Immunoblots were probed with the indicated antibodies. (**D**) Change in FRET-ratio after NCS addition depends on ATM. Time-lapse sequence (left) or quantification of 1/FRET (right) of U2OS cells expressing H2B-ATKAR. Time point 0 indicates addition of 20 nM NCS and time point 1 indicates addition of KU60019 (10 uM, ATMi), VE821 (1 uM, ATRi), or NU7026 (10 uM, DNAPKi). Graph shows average and SD of at least 15 cells. (**E**) GFP pull-down from U2OS cells expressing ATKAR wild-type (Wt) or alanine mutant (Ala) treated with NCS (10 nM) in presence or absence of ATM inhibitor. Immunoblot analysis shows the phosphorylation of ATKAR is on the expected target site residue and is ATM-dependent upon NCS treatment. (**F**) Quantification of 1/FRET of MCF-7 and RPE cells expressing H2B-ATKAR treated with the indicated concentrations of NCS. Graph shows average of at least 10 cells per condition.

**Figure 2–figure supplement.**
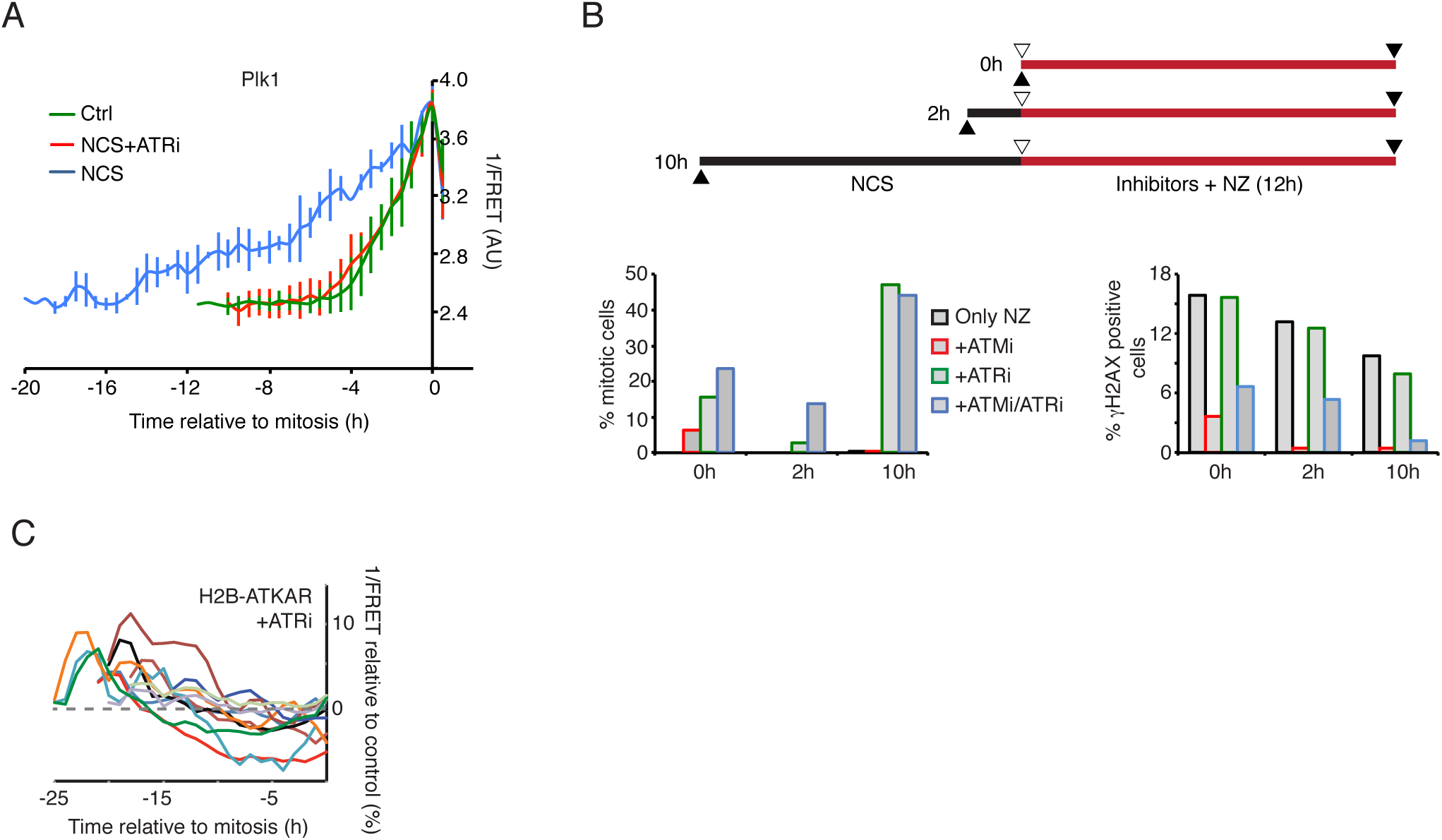
ATM inhibits Plk1 during the early phases of a DDR. (**A**) ATR inhibits Plk1 activity during checkpoint recovery in RPE cells. RPE cells expressing a Plk1 FRET-probe were transfected with p53 siRNA. NCS (8 nM) and ATRi (1 uM) were added as indicated. Graph shows average and SD of 15 cells (ATRi and Ctrl) or 2 cells (NCS; spontaneous recovery) synchronized *in silico* in mitosis. (**B**) Synergistic effect of ATM and ATR inhibition early after NCS. U2OS cells were treated with NCS (1 nM) for 0, 2, and 10 h, and subsequently incubated for 12 h with nocodazole and inhibitors as indicated. Cells were fixed, stained for pS10-histone H3 and γH2AX and analyzed by FACS. (**C**) H2B-ATKAR is dephosphorylated before cells enter mitosis in presence of ATR inhibitor. Quantification of 1/FRET of U2OS cells expressing H2B-ATKAR after treatment with VE821 (1 uM, 30 min) and NCS (2 nM). Each line represents a single cell that is synchronized in mitosis *in silico*. The FRET-ratio change of each cell relative to the FRET-ratio before NCS addition is shown.

**Figure 3–figure supplement 1.**
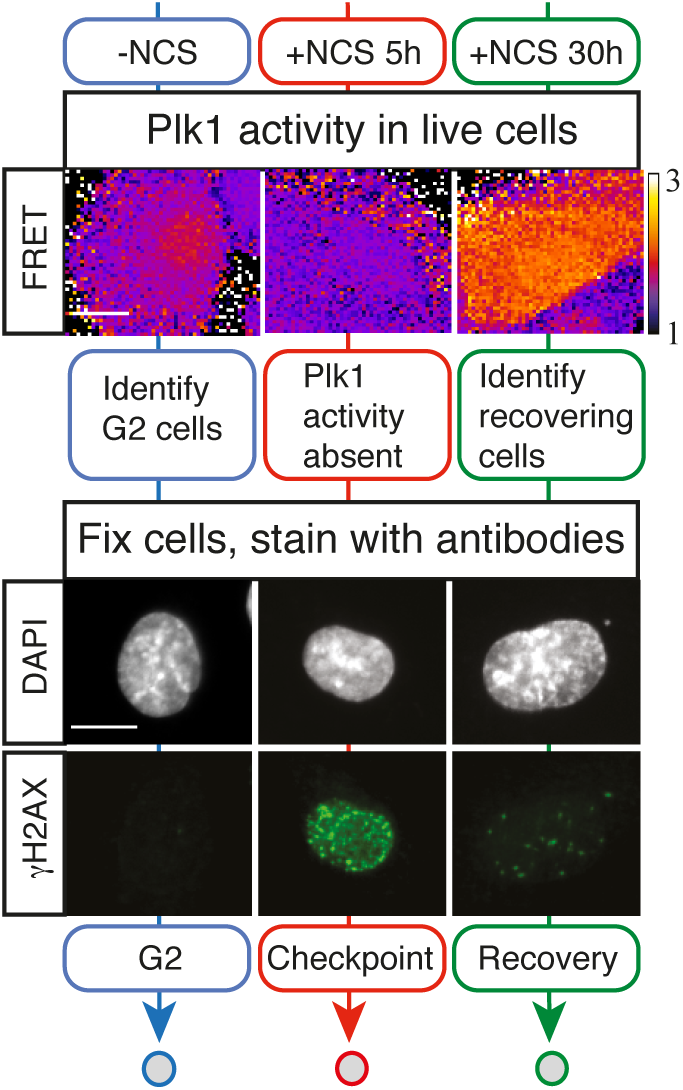
As opposed to H2B-ATKAR phosphorylation, ATM activity is sustained after Plk1 activation. Example of approach described in Figure 3A. Representative images of live cells depicting Plk1 activity and the same cells fixed and stained for γH2AX and DAPI are shown. Mock treated cells (blue), 5 h NCS (red) or 30 h NCS (green). Scale bar: 15um.

**Figure 3–figure supplement 2.**
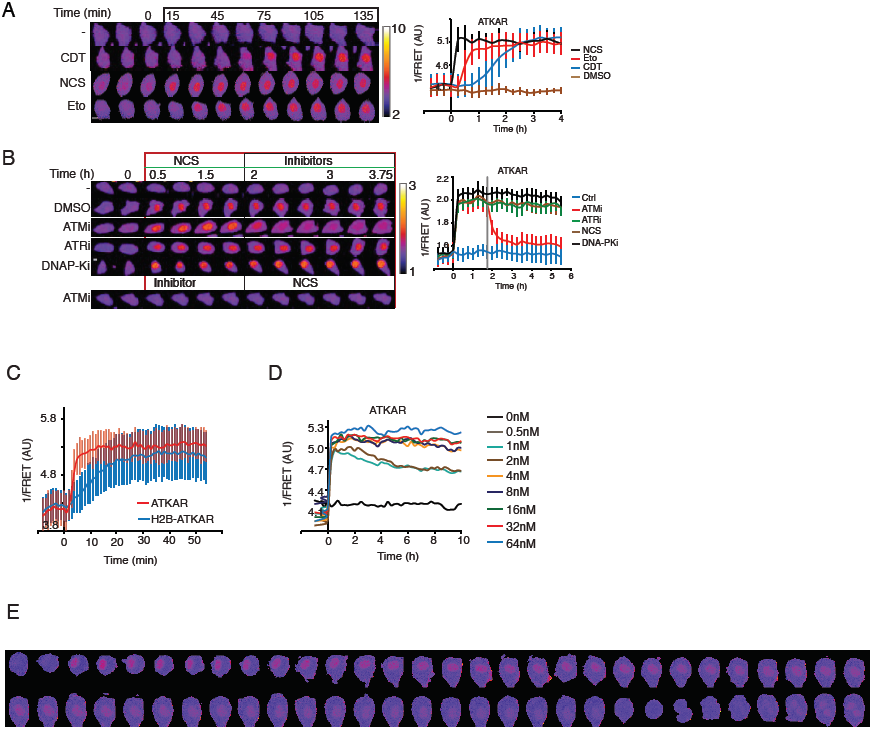
ATKAR and H2B-ATKAR detect different forms of ATM activity. (**A**) Kinetics of ATKAR 1/FRET change after treatment with Etoposide (Eto), Neocarzinostatin (NCS) or cytolethal distending toxin (CDT). Time-lapse sequence (left) or quantification of 1/FRET (right) of U2OS cells expressing H2B-ATKAR. Graph shows average and SD of at least 15 cells followed over time. Time point 0 indicates addition of drugs. (**B**) Change in FRET-ratio after NCS addition depends on ATM. Time-lapse sequence (left) or quantification of 1/FRET (right) of U2OS cells expressing ATKAR. Time point 0 indicates addition of 5 nM NCS and time point 1 indicates addition of KU60019 (10 uM, ATMi), VE821 (1 uM, ATRi), or NU7026 (10 uM, DNAPKi). Graph shows average and SD of at least 7 cells. (**C**) Quantification of 1/FRET of a mixed population of H2B-ATKAR and ATKAR expressing U2OS cells after addition of NCS (5 nM). H2B-ATKAR and ATKAR expressing cells were identified by the localization pattern of the expressed constructs. Graph shows average and SD of median pixel value of at least 7 cells. Time point 0 indicates addition of NCS. (**D**) Quantification of 1/FRET of U2OS cells expressing ATKAR, treated with the indicated NCS concentrations. Graph shows average of at least 10 cells per condition and is related to Figure 1A. **(E)** Example of ATKAR 1/FRET during checkpoint recovery. U2OS cells expressing ATKAR were treated with 1 nM NCS between time point 2 and 3 and the cell was followed over time. Note that ATKAR 1/FRET is sustained until mitotic entry and re-appears after mitosis. Time between images 30 min.

**Figure 4–figure supplement.**
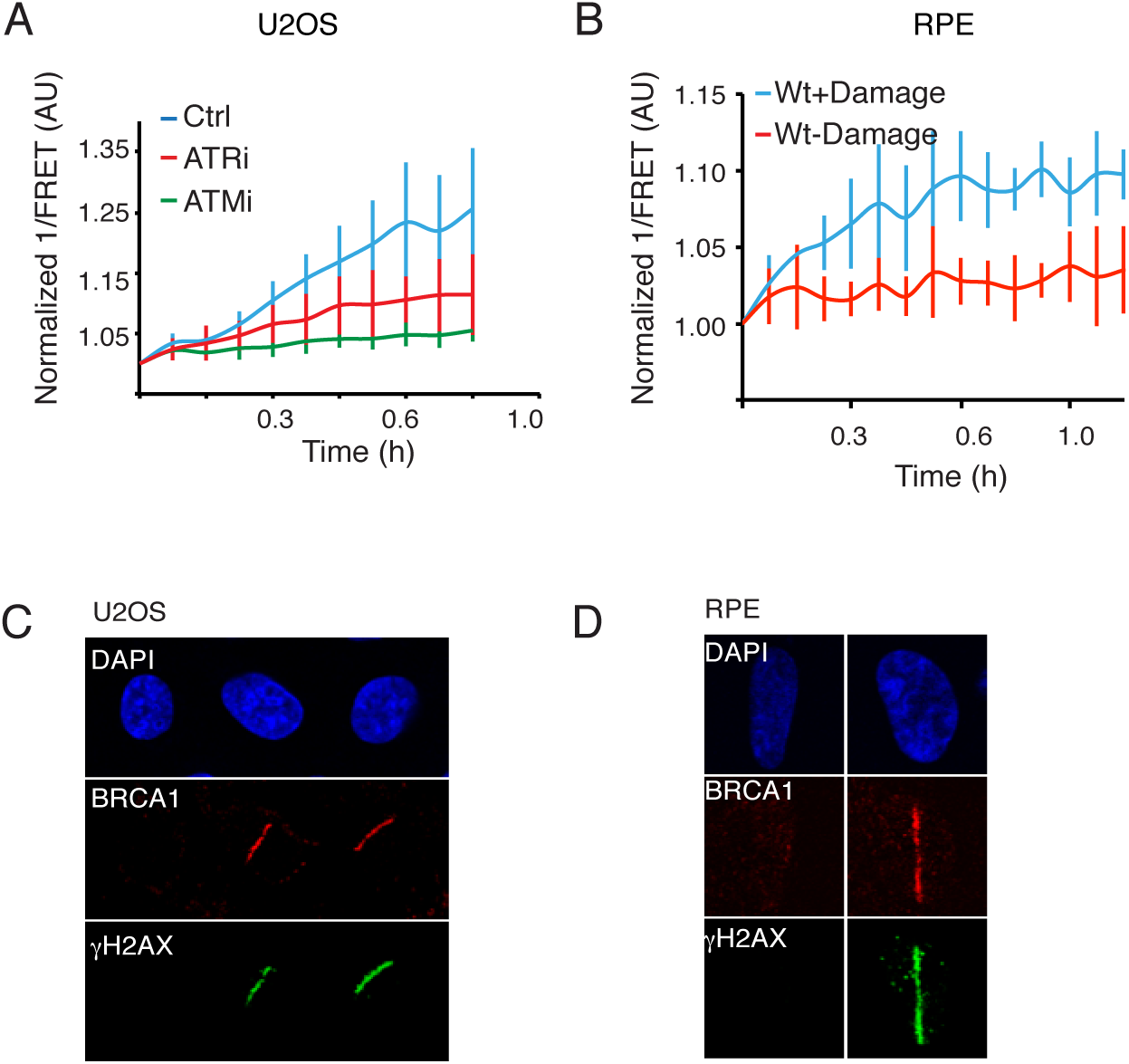
H2B-ATKAR detects spread of ATM activity over chromatin. (**A**) H2B-ATKAR detects both ATM and ATR activity after laser micro-irradiation. Quantification of spread of H2B-ATKAR FRET-change after laser microirradiation in U2OS cells in the presence of ATM or ATR inhibitors. Inhibitors were added 30 min before laser microirradiation. Graph shows average and standard deviation of at least 5 cells. Measurements were performed as in (Figure 4B). (**B**) Quantification of spread of H2B-ATKAR FRET-change after laser micro-irradiation in RPE cells. Measurements were performed distal to the laser-micro-irradiated area. Graph shows average and SD of at least 6 cells per condition, performed as in Figure 4B. (**C, D**) DSBs are restricted to the laser microirradiated area. γH2AX and BRCA1 remain in laser microirradiated area in U2OS (D) and RPE cells (E). Images show immunofluorescence stainings with indicated antibodies in laser microirradiated and neighbouring non-irradiated cells.

**Figure 7–figure supplement.**
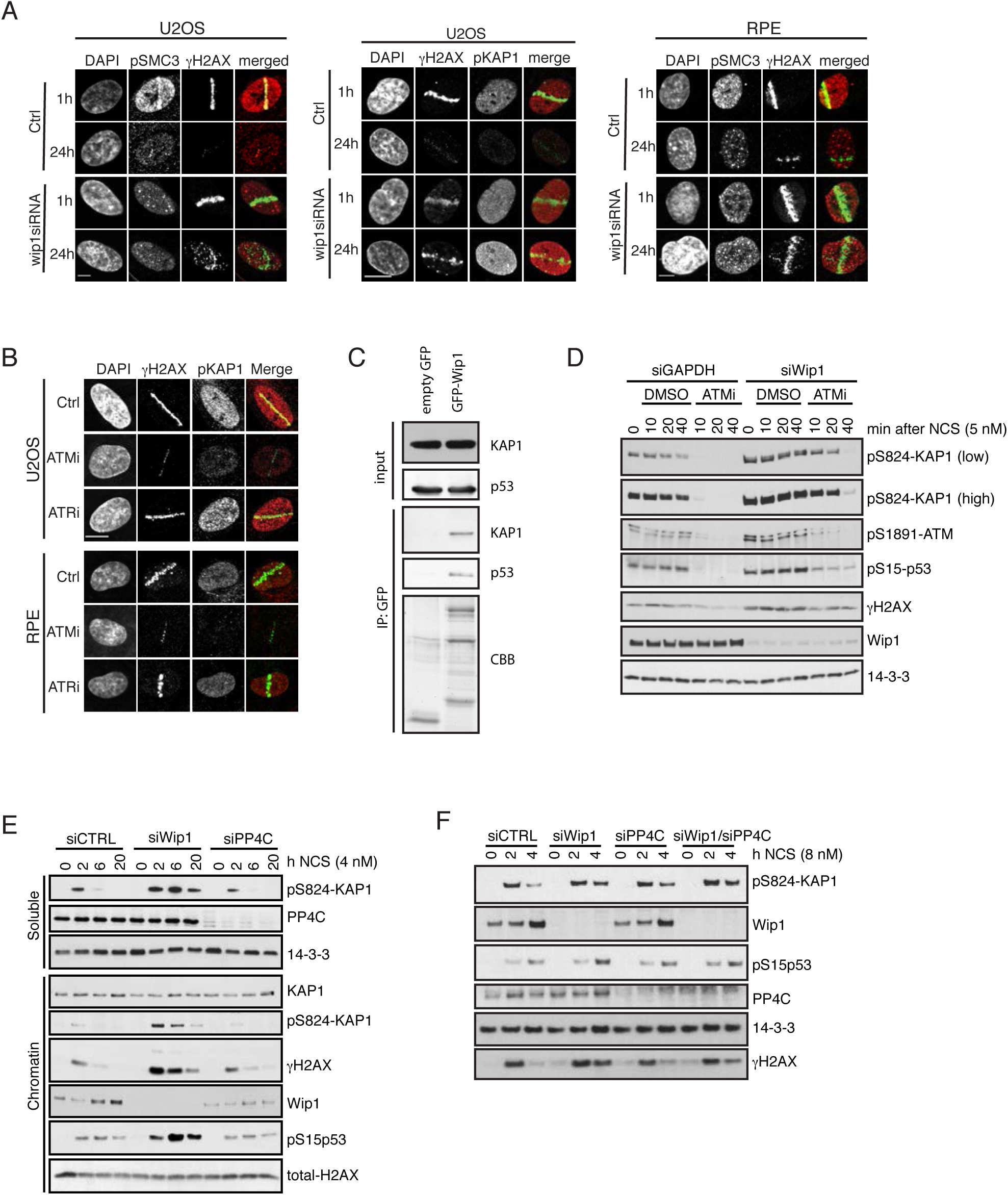
Wip1 dephosphorylates Kap1 pS824. **(A)** U2OS or RPE cells were transfected with Wip1 siRNA, microirradiated, and stained after 1 or 24 h with the indicated antibodies. **(B)** U2OS and RPE cells were microirradiated in the presence of indicated inhibitors. After 1 h, cells were stained with the indicated antibodies. **(C)** EGFP or EGFP-Wip1 was immunoprecipitated from HEK293 cells using GFP-Trap. Endogenous KAP1 and p53 were probed with antibodies. **(D)** U2OS cells transfected with GAPDH or Wip1 siRNA were treated with NCS in combination with DMSO or ATMi for indicated times. Whole cell lysates were probed with indicated antibodies. **(E)** U2OS cells transfected with GAPDH, Wip1 or PP4C siRNA were treated with NCS for indicated times. Soluble and chromatin fractions were probed with indicated antibodies. **(F)** RPE cells transfected with GAPDH, Wip1 or PP4C siRNA were treated with NCS for indicated times. Whole cell lysates were probed with indicated antibodies.

